# CellDiffusion: a generative model to annotate single-cell and spatial RNA-seq using bulk references

**DOI:** 10.1101/2025.10.27.684671

**Authors:** Xiaochen Zhang, Jiadong Mao, Kim-Anh Lê Cao

**Affiliations:** Melbourne Integrative Genomics, School of Mathematics and Statistics, The University of Melbourne, Australia

## Abstract

Annotating single-cell and spatial RNA-seq data can be greatly enhanced by leveraging bulk RNA-seq, which remains a cost-effective and well-established benchmark for characterising transcriptional activity in immune cell populations. However, a major technical hurdle lies in the contrasting properties of these data types: single-cell and spatial data are inherently sparse due to its cell-level sampling scheme, leading to much lower sequencing depth compared to bulk RNA-seq.

We developed CellDiffusion, a generative machine learning (ML) tool that bridges this gap. CellDiffusion generates realistic virtual cells to augment the sparse single-cell and spatial data, improving signals and the representation of rare cell types. The augmented data are more comparable to bulk references, increasing the accuracy of cell type annotation using bulk references and automated ML classifiers.

We benchmarked CellDiffusion on single-cell and spatial datasets from human peripheral blood samples, white adipose tissues, and breast tumours. Our method significantly outperforms state-of-the-art methods such as SingleR, Seurat, and scVI. In addition, CellDiffusion provides critical biological insights, including the identification of novel cell subtypes and their function during cell state transition; the discovery of new marker genes for tissue-resident immune cells, revealing their functional shifts in myeloid populations; and the accurate characterisation of cell subtypes in spatial transcriptomics to decipher tumour microenvironment.

## 1 Introduction

The landscape of transcriptomic research has been fundamentally altered by technologies that profile gene expression at single-cell resolution [1]. Single-cell RNA sequencing (scRNA-seq) enables the high-throughput dissection of complex tissues into their constituent cell types, providing critical insights into cellular heterogeneity, rare cell populations, and pathological states [2]. Complementing this, image-based spatial transcriptomics overcomes the loss of positional information inherent in dissociated-cell methods [3]. By co-registering transcriptomic data with spatial coordinates, these techniques allow for the direct visualisation of gene expression patterns within the native tissue architecture, achieving true single-cell resolution upon computational cell segmentation [4]. Their application offers a comprehensive view of biological processes, from characterising the tumour microenvironment to identifying spatially resolved biomarkers [5].

Realising the full potential of this high-dimensional single-cell resolution data hinges on a critical subsequent step: assigning a precise biological identity to every cell [6]. This process, known as cell type annotation, is fundamental to extracting meaningful insights and has become the essential framework for characterizing tissue composition, elucidating disease mechanisms, and understanding cell-specific responses to therapies [7]. The exponential growth in dataset size and complexity has, in turn, driven a necessary shift from laborious manual annotation to scalable, automated computational approaches [8].

Currently, automated cell type annotation strategies predominantly rely on mapping query datasets to scRNA-seq reference atlases [9]. However, the efficacy of this approach remains critically dependent on both the quality and biological relevance of the selected reference atlas [10]. Generating new, application specific scRNA-seq references is not only prohibitively expensive but also demands extensive manual cell type curation [11]. Furthermore, these references must be precisely tissue matched to the query data, as they generally do not perform well across different tissue types [12]. These limitations are further compounded in spatial transcriptomics applications, where studies have shown that annotation based on scRNA-seq atlas can produce inadequate results due to inherent technical constraints, including low signal-to-noise ratios that compromise accurate and robust cell type mapping [13].

An alternative strategy is to utilise the vast archives of bulk RNA sequencing data accumulated over decades. The high-throughput nature of bulk RNA-seq technology, coupled with numerous public repositories, renders this approach cost-effective and often eliminates the need for *de novo* reference atlas generation [14]. Furthermore, the higher signal-to-noise ratio and lower sparsity of bulk data make them inherently more robust for challenging tasks such as cross-tissue or cross-platform cell type annotation [15]. For years, these repositories have served as a cornerstone of clinical omics, creating a rich, biologically grounded standard coupled with meticulous clinical and pathological annotations from large patient cohorts [16]. Therefore, the integration of high-resolution single-cell data with established bulk RNA-seq resources offers a powerful analytical framework for aligning novel discoveries within existing clinical knowledge, effectively bridging the translational gap between laboratory findings and established clinical records [17, 18].

Various bioinformatics tools are available for the integration of single-cell resolution and bulk RNA-seq data. Prominent examples include SingleR, which uses correlation-based methods to provide accurate annotations, and Sincast, which enhances the sparse singlecell signal by imputation and pseudobulk aggregation to improve comparability with bulk data [19, 20]. Another strategy, used by methods such as Phi-space, involves projecting both data types into a common lower-dimensional space to enable integrated downstream analysis, including cell type annotation [21].

A key limitation of current methods for integrating single-cell and bulk RNA-seq data is their reliance on statistical methods alone [22]. These approaches never model the physical sampling process that causes the fundamental differences between single-cell and bulk RNAseq data. The single-cell sequencing experiment is inherently an undersampling process. Due to practical and cost constraints, only a fraction of cells are captured from the tissue with lower sequencing depth [1]. Recent advances in generative artificial intelligence (AI), particularly the development of diffusion models for creating high-quality data, present a powerful opportunity to overcome this limitation [23]. Here, we introduce CellDiffusion, a method that applies a generative model to simulate the transcriptomes of cells likely missed during experimental sampling. By augmenting the query single-cell and spatial RNA-seq dataset by using virtual cells, CellDiffusion makes it more comparable to comprehensive bulk RNA-seq atlases, thereby providing a more robust and cost-effective approach to cell type annotation. Our method aims to allow researchers to better utilise existing biological knowledge from bulk RNA-seq, potentially reducing the reliance on costly new reference atlases and improving the annotation of cell states that may be underrepresented in singlecell datasets.

## 2 Results

### 2.1 CellDiffusion to annotate cell types using bulk RNA-Seq references

#### Overview

The design of CellDiffusion is directly inspired by the data generation process in single-cell experiments. During tissue dissociation and cell capture, only a fraction of the cells from the original sample are successfully sequenced, resulting in an inherently incomplete representation of the tissue’s true cellular diversity. CellDiffusion addresses this undersampling issue by learning the underlying transcriptomic distributions from the observed cells to generate “virtual cells” with plausible transcriptional profiles of cells that were missed during the sequencing workflow. By aggregating virtual and observed cells, we create augmented cells that are directly comparable to bulk RNA-seq references while retaining local, single-cell resolution. This approach enables the use of robust classifiers for accurate cell type annotation. The complete conceptual model and modular workflow of CellDiffusion are presented in Figure 1.

**Fig 1:**
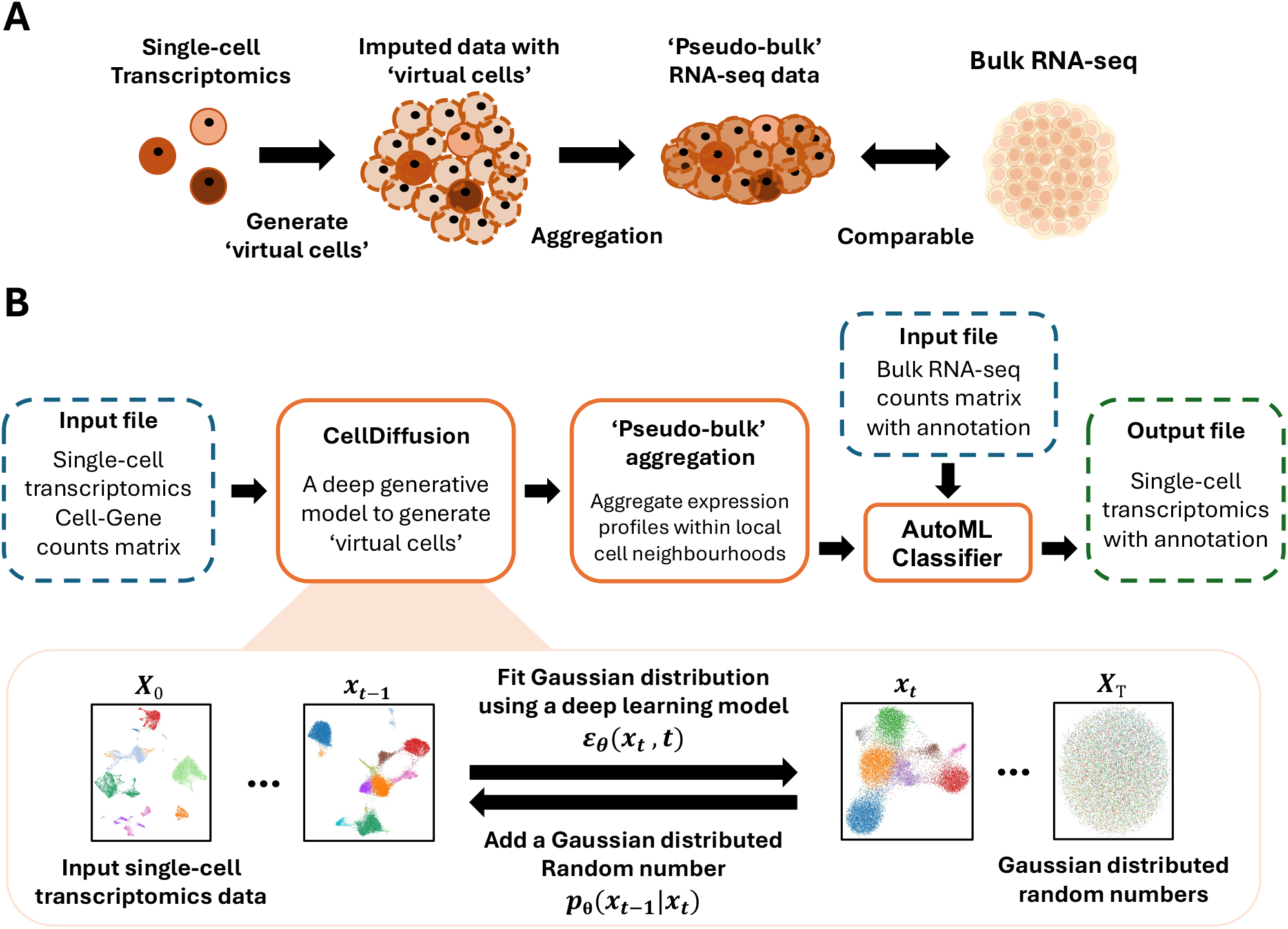
Overview of the CellDiffusion Framework. **A:** Conceptual model of CellDiffusion. Single-cell RNA-seq data represent an incomplete sample of the cellular diversity within tissues. CellDiffusion addresses this by computationally generating “virtual cells” to represent the unsampled portion of the population. **B:** The workflow starts with a core denoising diffusion probabilistic model (DDPM) that generates “virtual cells”. The DDPM learns to reverse a forward noising process, allowing the generation of new, realistic virtual cells from a random input. The generated cells faithfully capture the underlying data distribution learned from the real cell population. Then, a cell augmentation module pools the original and virtual cells to create profiles comparable to bulk RNA-seq data. Finally, an automated machine learning classifier annotates these profiles using an established bulk RNA-seq reference atlas.

#### A denoising diffusion generative model

The core of our framework is based on a denoising diffusion probabilistic model (DDPM) trained on single-cell and spatial RNA-seq data. By learning the underlying data distribution (Figure 1B), the model generates a large population of synthetic, biologically plausible “virtual cells”. This generative process enriches the dataset at a low computational cost, creating a more complete representation of the cellular landscape than initial, experimentally derived sample. Crucially, this approach is highly effective for imputing the transcriptomic profiles of rare or transient cell states, which are often missed or underrepresented due to experimental sampling limitations.

#### Cell augmentation module

To bridge the technical gap between single-cell and bulk sequencing modalities, this module pools small, similar groups of original and generated cells (typically around 15 cells) into local neighbourhoods (Figure 1B). By generating a unique augmented cell centred on each individual real cell, this method augments the sparse singlecell data, improving its signal quality to be comparable with robust bulk references. This cell-centric strategy is critical: it enables the use of informative bulk atlases for annotation while preserving the single-cell resolution of the data.

#### Automated machine learning classifier

For the final annotation step, we employ an automated machine learning (AutoML) classifier that operates on the high-quality augmented cells. This module systematically searches for the optimal classification model and hyperparameters for any given bulk reference dataset (Figure 1B). By automating model selection, this approach ensures robust and highly accurate annotations across diverse biological contexts and reference types, eliminating the need for laborious manual tuning. This design places the emphasis on the quality of the input signal rather than on a fixed choice of classifier, as the system automatically identifies a high-performing model when the biological signal is sufficient.

#### Validation and case studies

We demonstrate the efficacy and versatility of our framework through rigorous benchmarking and three distinct case studies. First, we quantitatively assess CellDiffusion’s performance against established annotation methods. We then showcase its ability to uncover novel biological insights through applications in profiling peripheral blood mononuclear cells, analysing white adipose tissue, and interrogating the tumour microenvironment with spatial transcriptomics.

### 2.2 Benchmark study

To systematically evaluate CellDiffusion’s performance, we conducted a comprehensive benchmarking against four widely used annotation methods: SingleR, Seurat, CHETAH, and scVI (scANVI), across three distinct single-cell resolution datasets spanning different tissue types, technologies, and biological contexts [19, 22, 24, 25].

Uniform Manifold Approximation and Projection (UMAP) visualisation of cell type annotations (Figure 2A) demonstrates CellDiffusion’s superior ability to recapitulate ground truth cell type distributions when using bulk RNA-seq references. While SingleR and Seurat also accommodated bulk references, they generated less coherent clustering patterns [19, 22]. In particular, CHETAH and scVI, designed primarily for scRNA-seq references, failed to generate meaningful annotations when provided with bulk RNA-seq references, highlighting the technical challenge that CellDiffusion effectively addresses [24, 25].

**Fig 2:**
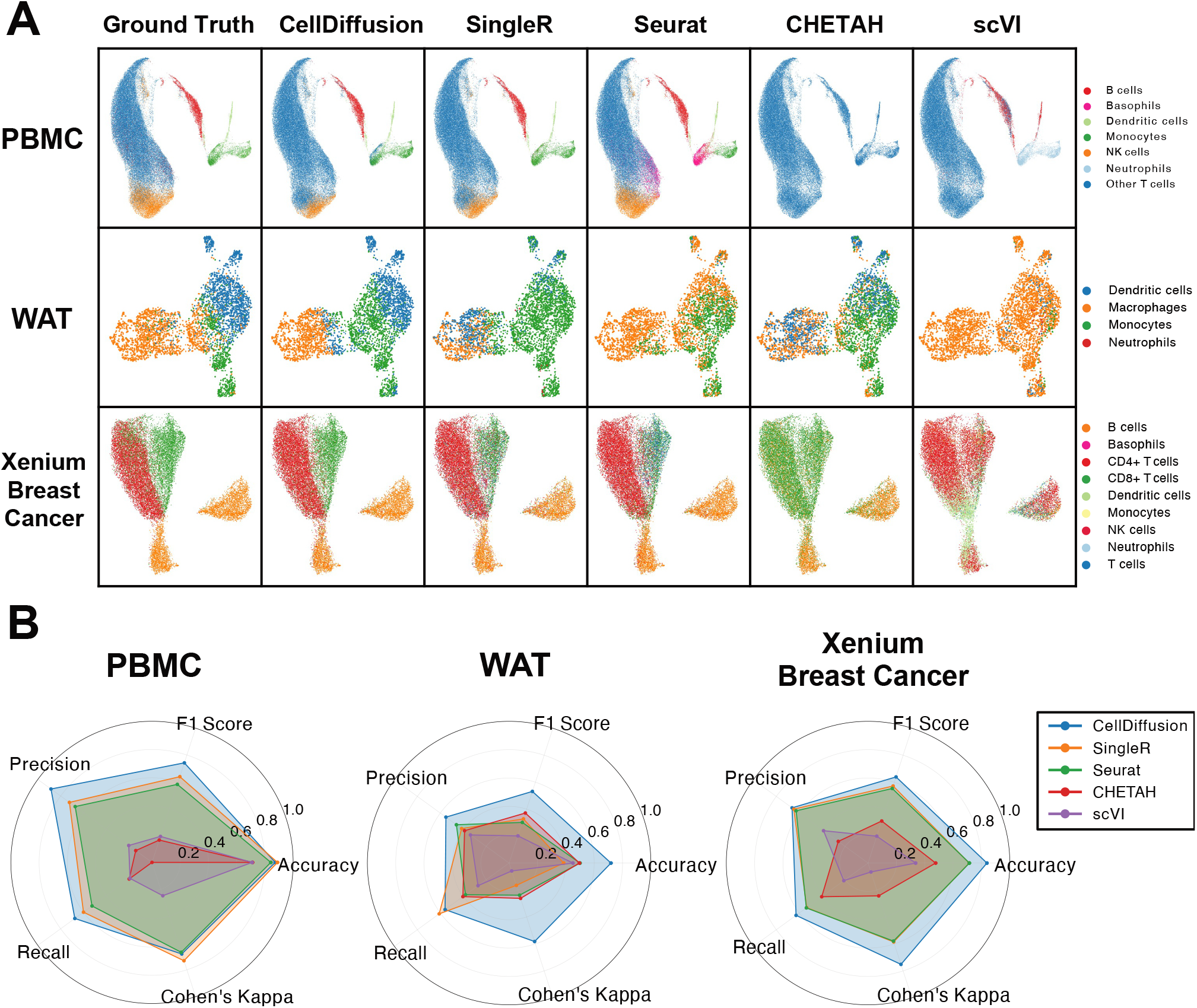
Benchmarking CellDiffusion against established cell type annotation methods. **A:** UMAP visualisations of cell type annotations on three distinct datasets, using a bulk RNA-seq reference. CellDiffusion’s annotations closely recapitulate the ground truth distribution, generating coherent and well-separated cell clusters. In contrast, while SingleR and Seurat can process the bulk reference, they produce less distinct clustering patterns. Methods not designed for this task, CHETAH and scVI, fail to generate meaningful annotations. **B:** Radar plots comparing performance across five key metrics for the three datasets. CellDiffusion outperforms competing methods across the majority of evaluations. The larger area covered by CellDiffusion in each plot signifies its superior overall performance across all tested datasets.

Quantitative performance metrics further confirm CellDiffusion’s advantages (Figure 2B). Across the three datasets, radar plots synthesising key metrics (accuracy, precision, recall, F1 score, and Cohen’s Kappa) demonstrate that CellDiffusion consistently outperforms alternative methods. While SingleR showed slightly higher performance in terms of accuracy and Cohen’s Kappa on the PBMC 68k dataset and for recall on the adipose dataset, CellDiffusion achieved the highest overall score across the vast majority of evaluations. CellDiffusion’s superior performance was particularly pronounced when annotating rare cell populations, where traditional correlation-based (e.g. SingleR) and projection-based methods (e.g. Seurat) often struggle due to limited signal-to-noise ratios.

In summary, our benchmarking results established CellDiffusion as a competitive method for single-cell and spatial RNA-seq annotation using bulk RNA-seq references. Across diverse datasets, CellDiffusion consistently delivered more accurate and coherent cell type assignments than leading tools. Its key advantage lies in the robust identification of rare cell types, a task where conventional methods often fall short.

### 2.3 Identification of novel monocyte subtypes in human PBMC scRNA-seq data using CellDiffusion

We illustrate the identification of novel cell subtypes and cellular states with CellDiffusion to identify on a publicly available scRNA-seq dataset of human peripheral blood mononuclear cells (PBMCs) [26]. The complex and well-characterised nature of this immune cell population provides a compelling biological context to demonstrate the discovery power of our method.

While CellDiffusion successfully annotated all major immune lineages with high accuracy (Figure 3A), its superior resolution became evident in the analysis of monocytes. Focusing on this key myeloid population, CellDiffusion resolved three distinct subtypes: classical, intermediate, and non classical monocytes (Figure 3B1). This result provides a more granular view than standard analysis pipelines; for comparison, the widely used Azimuth reference mapped the same data to only two subtypes, classical and non classical monocytes, failing to distinguish the intermediate population (Figure 3B2).

**Fig 3:**
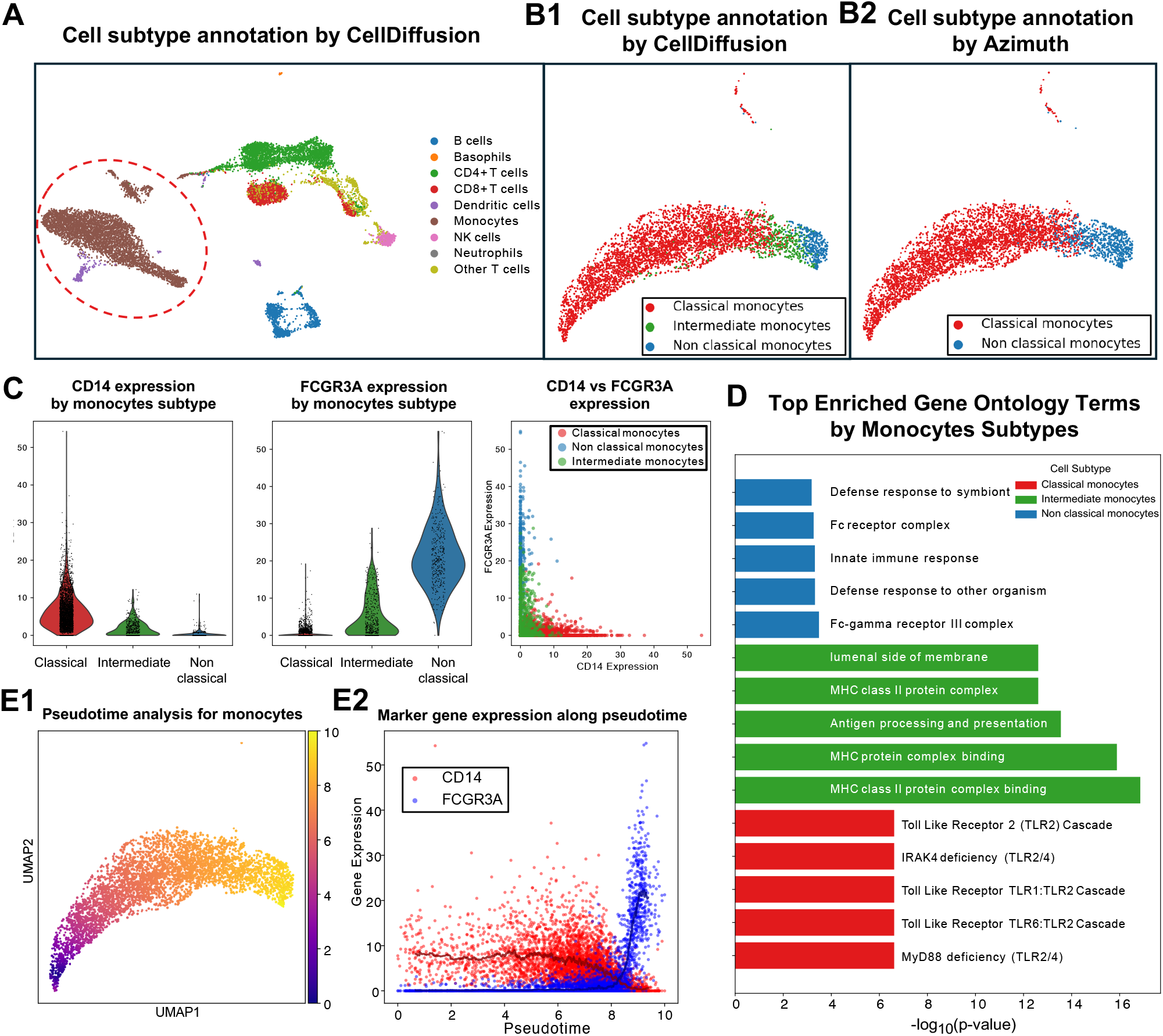
CellDiffusion identifies and characterises a distinct intermediate monocyte subtype in human PBMCs. **A:** UMAP visualisation of major cell types annotated by CellDiffusion in a public human PBMC dataset. Monocytes (highlighted) were isolated for detailed subtype analysis. **B:** Comparative UMAPs of monocyte subtypes. B1: CellDiffusion, using a bulk RNA-seq reference, resolves three populations: classical, intermediate, and nonclassical. B2: Azimuth, using a scRNA-seq reference, identifies only two populations: classical and non classical monocytes. **C:** Violin plots showing normalised expression of canonical markers CD14 and FCGR3A across the three monocyte subtypes, confirming the distinct expression profile of the intermediate population. **D:** Functional enrichment analysis of marker genes for intermediate monocytes. The top enriched terms are associated with antigen processing and presentation via MHC class II, suggesting a specialised immunological function. **E:** Pseudotime analysis models the differentiation trajectory from classical to nonclassical monocytes. E1: The trajectory clearly positions the intermediate population as a transitional state. E2: Gene expression dynamics along pseudotime show a decrease in CD14 and a concurrent increase in FCGR3A, consistent with this differentiation path.

The intermediate monocyte subtype identified by CellDiffusion was characterised by the simultaneous expression of CD14 and FCGR3A, as visualised in the UMAP plot of marker gene expression (Figure 3C). This unique expression pattern distinguishes intermediate monocytes from classical and non classical monocytes, suggesting that they represent a distinct cell population.

Importantly, this intermediate population represents more than a transitional state: this population has its own unique functional role. The enrichment analysis revealed that intermediate monocytes were significantly enriched for genes involved in the MHC class II antigen presentation pathways (Figure 3D). These pathways are fundamental to adaptive immunity, enabling specialised cells to present foreign antigens to helper T cells, which in turn orchestrate a targeted and lasting immune response [27]. Our analysis, therefore, pinpoints intermediate monocytes as professional antigen presenting cells (APCs), whose main function is to activate the adaptive immune system. This role is functionally distinct from the primary phagocytic duty of classical monocytes and the patrolling surveillance of non classical monocytes [28]. This conclusion is strongly supported by previous work that has identified intermediate monocytes as the most potent antigen-presenting subset in their constitutive state [29].

To place this functionally distinct subtype within its developmental context, we performed pseudotime trajectory analysis. The results inferred a continuous differentiation path from classical to non classical monocytes, with the intermediate cells positioned as a transitional state between them (Figure 3E). The smooth decrease in CD14 expression and the concurrent increase in FCGR3A expression along this trajectory provide strong evidence for a differentiation continuum.

In summary, by integrating its fine-grained annotations with downstream analyses, our study provides a more nuanced view of cellular differentiation, demonstrating that transitional cells within a continuum can be functionally specialized. These subtle cell states are often overlooked by standard scRNA-seq pipelines, which typically lack the robust highresolution reference data needed to identify them. For instance, by leveraging bulk RNA-seq references in our analysis of PBMCs, CellDiffusion identified a functionally specialized intermediate monocyte population missed by other methods. This discovery underscores the tool’s ability to refine our understanding of cellular heterogeneity and the dynamic roles of transitional cells.

### 2.4 CellDiffusion reveals novel tissue-specific markers for immune cells in adipose tissue

We evaluated CellDiffusion’s ability to characterise tissue-specific cellular features on a human white adipose tissue scRNA-seq dataset (Figure 4A1) [30]. This tissue is an ideal test case as it contains diverse populations of resident immune cells, which adapt to the local microenvironment and often diverge phenotypically from circulating immune cells [31].

**Fig 4:**
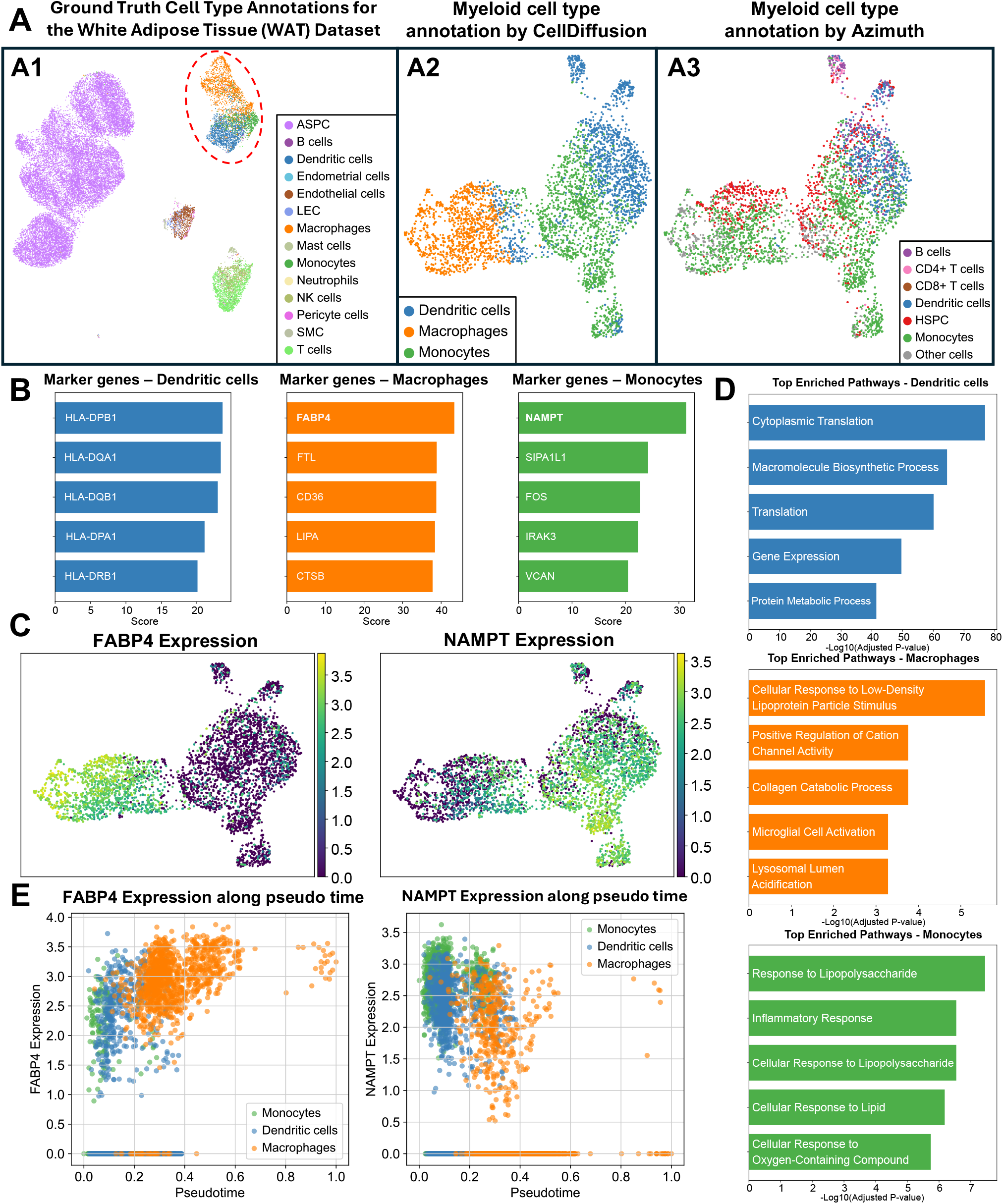
CellDiffusion accurately annotates tissue-resident immune cells and identifies novel markers in adipose tissue. **A:** UMAP projections of adipose tissue scRNA-seq data [30], focusing on myeloid cells. A1: Global view of all cell types with myeloid populations highlighted. A2: Cell type annotations of the isolated myeloid subset by CellDiffusion (using a bulk RNA-seq reference) A3: Annotations of the myeloid subset by Azimuth (using a scRNA-seq reference).**B:** Bar plot of marker genes from differential expression analysis of CellDiffusion annotations. Analysis confirms known markers and identifies novel, highly significant markers such as FABP4 and NAMPT. **C:** Gene Ontology (GO) enrichment analysis for the three cell type populations. Enriched terms are consistent with known functions and highlight adipose-specific roles. **D:** UMAP feature plots showing distinct expression of the novel markers FABP4 (macrophages) and NAMPT (monocytes). **E:** Pseudotime trajectory analysis of monocyte to macrophage differentiation, revealing opposing expression dynamics where FABP4 increases and NAMPT decreases along the differentiation path.

Using a general bulk RNA-seq reference of myeloid cell lines, CellDiffusion accurately annotated the major adipose-resident myeloid cell types, showing high concordance with the original study’s manual annotations (Figure 4A2) [30]. This performance notably exceeded that of a standard Seurat workflow with the Azimuth scRNA-seq reference (Figure 4A3). Importantly, neither the bulk reference nor Azimuth contained adipose-tissue-specific myeloid cells, yet Azimuth failed to correctly resolve the monocyte and macrophage populations. This discrepancy underscores a key challenge: tissue-resident myeloid cells diverge significantly from their counterparts in reference atlases due to tissue-specific adaptations, and CellDiffusion’s approach appears more robust to this variation.

In addition, the robust annotations provided by CellDiffusion enabled a detailed exploration of the adipose-resident myeloid scRNA-seq data. Differential expression analysis of the annotated populations first confirmed established markers, including HLA family genes in dendritic cells, CD36 in macrophages, and FOS in monocytes (Figure 4B) [32–34]. Beyond this validation, the analysis revealed FABP4 and NAMPT as highly significant and specific markers for adipose-resident macrophages and monocytes, respectively. Their expression was tightly restricted to these populations (Figure 4C), reflecting functional adaptation to the tissue microenvironment. FABP4 is a key protein in lipid metabolism, likely involved in processing lipids from surrounding adipocytes [35], while NAMPT is a rate-limiting enzyme in NAD^+^ biosynthesis, pointing to specialised metabolic programming [36]. These findings are consistent with existing literature, which has demonstrated the function of FABP4 in adipose-resident macrophages [37] and has established the importance of NAMPT for both monocyte function and adipose tissue biology [38, 39].

This theme of metabolic adaptation was reinforced by gene ontology enrichment analysis. Both macrophages and monocytes showed significant enrichment for pathways related to lipid handling and metabolic responses, such as “cellular response to low-density lipoprotein particle stimulus” and “response to lipopolysaccharide” (Figure 4D). These functional profiles contrast with the primarily inflammatory signatures of circulating immune cells, highlighting a shift towards metabolic roles within the adipose tissue. To investigate the dynamics of these markers during monocyte to macrophage differentiation, we performed a pseudotime trajectory analysis. This analysis revealed opposing expression patterns: FABP4 expression increased along the differentiation trajectory, while NAMPT expression decreased (Figure 4E). This dynamic demonstrate the crucial role of FABP4 for macrophages and NAMPT for monocytes. It also suggests that these genes may play distinct regulatory roles during macrophage maturation within the adipose tissue microenvironment.

This case study highlights CellDiffusion’s unique strength in leveraging general bulk RNA-seq references to accurately resolve tissue-specific cell populations that are challenging for conventional scRNA-seq atlases. The precision of CellDiffusion’s annotations enabled both the validation of known markers and the discovery of novel, tissue-adapted marker genes like FABP4 and NAMPT. By providing a more accurate cellular map of the tissue, CellDiffusion directly facilitated deeper biological insights into the metabolic reprogramming of resident immune cells. Ultimately, this demonstrates CellDiffusion’s value as a powerful discovery tool capable of revealing nuanced, tissue-specific biology that might otherwise be obscured when relying on canonical cell references [40].

### 2.5 CellDiffusion resolves the spatial architecture of the tumour microenvironment in breast cancer

We applied CellDiffusion to a Xenium spatial transcriptomics dataset from breast cancer samples to demonstrate its utility on image-based spatial transcriptomics data. We focused on resolving the immune landscape of the tumour microenvironment (TME), where the spatial arrangement of immune cells is crucial for understanding anti-tumour responses.

Using a bulk RNA-seq reference of immune cells, CellDiffusion successfully annotated the major immune populations within the TME. The resulting spatial cell type maps revealed distinct cellular clusters (Figure 5A), clearly identifying populations of CD4^+^ T cells, CD8^+^ T cells, and B cells in proximity to the tumour region (Figure 5B).

**Fig 5:**
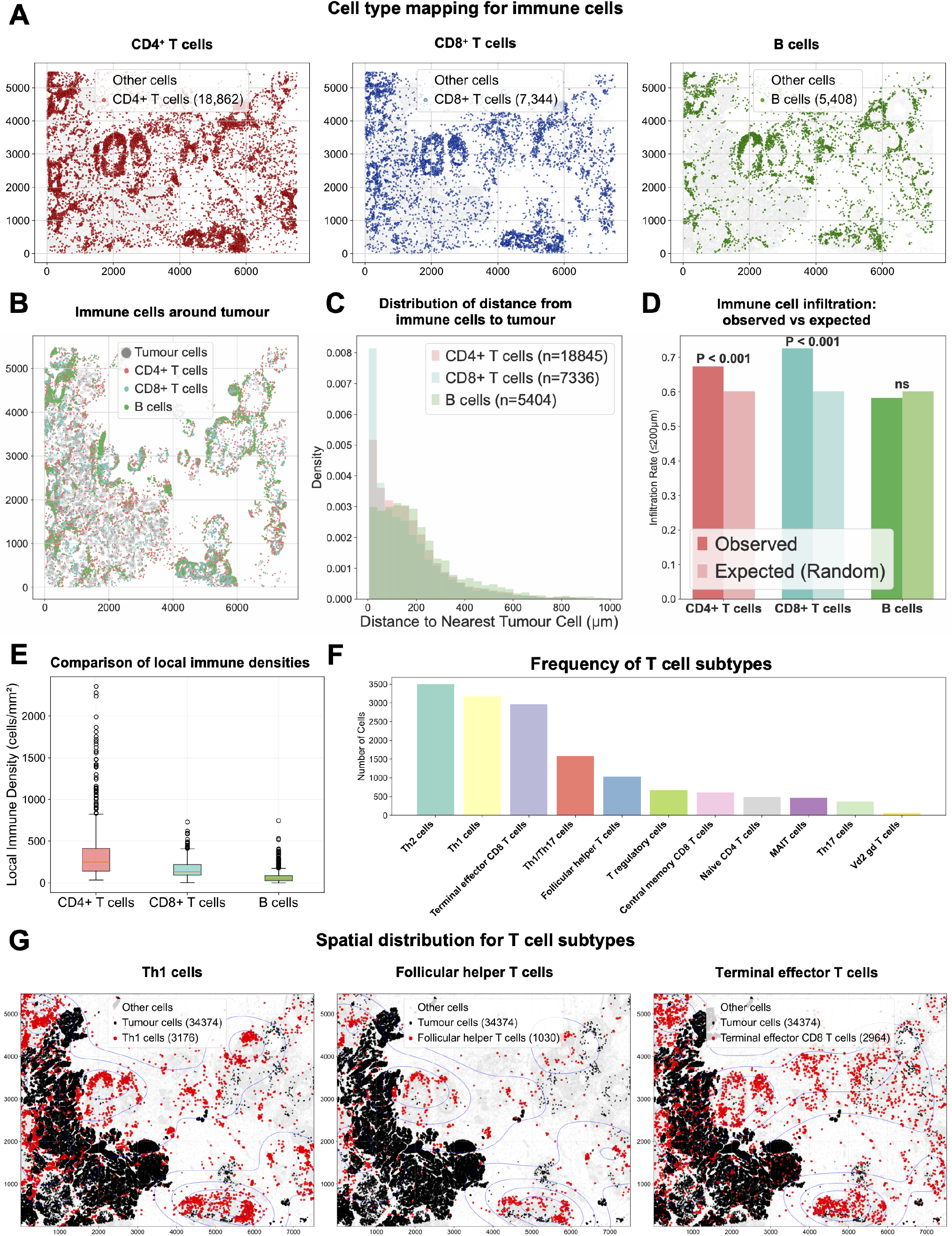
CellDiffusion annotate cell type for image-based spatial transcriptomics by using bulk RNA-seq reference. **A:** Pseudo-colour from cell type annotation of spatial transcriptomics data for CD4^+^ T cells, CD8^+^ T cells, B cells from breast cancer tissue by using CellDiffusion framework with bulk RNA-seq reference, showing distinct clusters corresponding to major cell populations in the tumour microenvironment. **B:** Immune cell selected near tumour region for downstream analysis. **C:** The distribution of the distance between the annotated cell types and the tumour region, showing that CD4^+^ T cells are the most abundant in the tumour region. CD4^+^ T cells are the second most abundant, and B cells are the least abundant. **D:** Immune cells enrichment analyses show that CD4^+^ T cells and CD8^+^ T cells are significantly enriched around the tumour, but B cells are not. **E:** Local Immune cell density shows that CD4^+^ T cells have the highest cell density around the tumour and CD8^+^ T cells are the second highest, while B cells have the lowest cell density. **F:** The cell subtype distribution of T cells, showing that Th1, Th2 and terminal effector T cells are the most abundant T cells with the highest proportion, while Tfh cells have a lower proportion than we expect. **G:** Spatial map highlighting the distinct distributions of key T cell subtypes, including the peri-tumoural clustering of Th1 cells, the high local density of Terminal Effector CD8^+^ T cells, and the exclusion of Tfh cells from the tumour interior.

We next quantified the spatial relationships between the annotated immune populations and the tumour. This analysis revealed a clear spatial heterogeneity of lymphocytes: CD4^+^ T cells were the most abundant population within the tumour region, followed by CD8^+^ T cells, whereas B cells were sparse (Figure 5C). Consistent with this, both CD4^+^ and CD8^+^ T cells were significantly enriched and exhibited the highest local density near the tumour, in stark contrast to B cells, which showed no significant enrichment and the lowest density (Figures 5D-E).

Our analysis revealed a multi-faceted immune dysfunction within the tumor microenvironment. The most striking feature was a compromised cytotoxic response, characterized by a large population of terminally exhausted CD8^+^ T cells (Figure 5F), indicating T cell exhaustion and a diminished anti-tumour effort [41, 42]. This weakened attack was compounded by a failure to mount an effective B cell response. We observed a significant scarcity of T follicular helper (Tfh) cells, which are essential for activating and sustaining B cell activity [43]. Spatially, these Tfh cells were almost entirely excluded from the tumor core, providing a clear mechanistic explanation for the observed lack of B cell infiltration 5G). Finally, the T helper landscape showed a slight polarization towards a Th2 phenotype, a known mechanism of tumor immune evasion [44].

This case study highlights CellDiffusion’s power to resolve complex cellular architecture from highly sparse, image-based spatial transcriptomics data. By resolving these nuanced T cell subtypes and their spatial organization, our method overcomes significant data limitations. This enables the generation of functional hypotheses that directly link cellular organization to disease mechanisms, effectively bridging the gap between existing bulk RNA-seq references and new spatial transcriptomics data.

## 3 Discussion

CellDiffusion is a deep learning framework designed to address a central challenge in transcriptomics: the integration of large, well-annotated bulk RNA-seq datasets with highresolution single-cell and spatial data. By generating virtual cells via a diffusion model, our approach helps bridge these modalities, mitigating some of the technical and distributional disparities that have traditionally hindered their integration. Our contribution allows researchers to utilise decades of existing biological knowledge from bulk RNA-seq, reducing the need for costly new reference atlases and providing access to a richer diversity of cellular states that may be underrepresented in single-cell datasets.

Our framework facilitates the elucidation of nuanced biological processes across diverse contexts. For example, we identified subtle, intermediate monocyte states in PBMCs. In adipose tissue, CellDiffusion’s ability to account for tissue specific expression patterns was essential for accurately characterising specialised immune cells. In the application to spatial transcriptomics, we highlighted CellDiffusion’s capacity to handle highly sparse data while preserving spatial context. Our case studies demonstrate that CellDiffusion is more than a cell annotation tool, but also a hypothesis-generating framework to reveal complex cellular identities and states.

While CellDiffusion shows promise, it is important to acknowledge its current limitations. Firstly, the accuracy of annotations is fundamentally dependent on the quality and comprehensiveness of the reference bulk RNA-seq atlas. An incomplete or biased reference may limit the ability to identify certain cell populations. Furthermore, the reliance on high-purity bulk references often generated via cell sorting currently tailors the method’s application primarily to immune cell types. Secondly, to make sure the generated virtual cells remain biologically accurate, our method currently uses a filtering step after generation. This process uses a k-NN graph to select the most realistic virtual cells. While effective, the necessity for this step indicates that the realism of the core generative model could be further improved. Finally, the training process for the diffusion model is computationally intensive and generally requires GPU acceleration, which may pose a challenge for users with limited computational resources, particularly when working with very large datasets.

This work paves the ways for future improvement, such as developing more efficient generative models to simultaneously enhance the quality of virtual cells and reduce running time. Creating models capable of predicting entire single-cell transcriptomic landscapes under specified biological conditions would represent a paradigm shift, enabling researchers to supplement laboratory work with powerful *in silico* experiments to accelerate discovery.

## 4 Methods

### 4.1 CellDiffusion generative model

CellDiffusion is a denoising diffusion probabilistic model (DDPM) [23] trained to learn the underlying data distribution of single-cell resolution transcriptomics and generate synthetic data, which we term “virtual-cells”. The model is versatile, capable of processing gene expression profiles from technologies like droplet-based scRNA-seq and image-based spatial transcriptomics.

The overall process involves three key stages: data preprocessing, model training, and virtual cell generation. First, raw gene expression data undergoes a standard preprocessing pipeline, including filtering, normalisation, and log transformation [45]. A set of highly variable genes is selected, and each cell’s expression vector is reshaped into a square matrix, *x*_0_, to be compatible with the model’s convolutional architecture.

Conceptually, the DDPM framework operates through a forward process, where Gaussian noise is incrementally added to the input data, and a learned reverse process. The core of CellDiffusion is a U-Net model [46] trained to reverse this noising process. Starting from pure noise, the trained model can iteratively denoise a sample to generate a new, synthetic gene expression matrix, 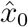. To account for inherent limitations in model performance, we employ a generate and filter strategy. Initially, a large surplus of virtual cells is generated. Subsequently, during the cell augmentation stage, a filtering step is implemented to select a high-quality subset of virtual cells, using information from the real cell population, before they are aggregated. This approach is computationally feasible due to the low cost of synthesising virtual cells.

A detailed description of the DDPM theoretical foundation, specific model architecture, training parameters, and the mathematical formulation for data generation is provided at Supplemental Material S1.

### 4.2 Cell augmentation

We used a pseudobulk aggregation strategy to bridge the gap between single-cell and bulk RNA-seq data. This method synthesizes bulk-like expression profiles from local cellular neighborhoods, which augments the signal for each cell while retaining the transcriptional heterogeneity inherent to the single-cell data.

The process begins by constructing a k-nearest neighbour (k-NN) graph on the combined dataset of original cells and the synthetic cells generated by the diffusion model. We utilise the scanpy package for this purpose, with the number of neighbours (*k*) set to a default of 15. The local community, *C*_*j*_, for a given cell *j* is defined as the set containing the cell itself and its *k*-nearest neighbours in the graph.

For each community, we generate a single augmented cell by averaging the gene expression vectors of all cells within that community. This aggregation step is defined by the formula:

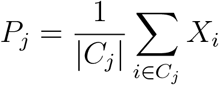

where *P*_*j*_ is the resulting augmented cell for community *j, X*_*i*_ is the expression vector of an individual cell *i* within that community, and |*C*_*j*_| is the total number of cells in the community (i.e., *k* + 1).

Finally, to ensure compatibility with real bulk RNA-seq data, the generated augmented cell are normalised to counts per million (CPM) and log-transformed. This neighbourhood-based aggregation effectively enhances the signal for distinct cellular states, including rare populations, and creates a well-founded dataset for direct comparison with bulk references.

### 4.3 Automated machine learning for cell type annotation

We employed an automated machine learning (AutoML) approach to classify our augmented cells using the bulk RNA-seq data as a reference. For this task, we utilised the Tree-based Pipeline Optimization Tool (TPOT, `version 0.12.2`), an AutoML framework that uses genetic programming to systematically discover the most effective machine learning pipeline [47]. TPOT was configured to search for an optimal classifier with a population size of 50 potential pipelines over 5 generations. This process automatically identified the best-performing model for assigning cell type labels to the augmented cells query data based on the patterns learned from the bulk reference.

### 4.4 Data and data preprocessing

We provide a comprehensive list of all datasets used in this study in Supplemental Material S2. We also specify the role of each dataset as either a query or a reference. We then detail the preprocessing pipelines for both data types. For query data, this includes the selection of highly variable genes, normalisation, log-transformation, and the crucial step of reshaping gene vectors into square matrices for compatibility with our model’s architecture. A parallel preprocessing workflow is described for the bulk RNA-seq reference data to ensure consistency.

### 4.5 Methods for benchmark and case studies

To validate our model, we conducted a rigorous benchmark study against several state-of-the-art annotation tools, including SingleR, Seurat, CHETAH, and scVI (scANVI). The Supplemental Material S3 details the specific parameters used for each method and defines the suite of performance metrics, such as the macro F1-score and Cohen’s Kappa, chosen for their robustness to class imbalance.

Furthermore, we outline the downstream analytical pipelines for our three biological case studies at Supplemental Material S4. These include methods for investigating cell sub-type heterogeneity and differentiation dynamics in monocyte and adipose tissue data (e.g., UMAP, differential gene expression, trajectory inference) and techniques for characterising the tumour microenvironment in spatial transcriptomics data (e.g., proximity analysis with permutation testing).

## Declarations

### Code availability

The CellDiffusion Python package is available on GitHub (https://github.com/ShiltonZhang/CellDiffusion), along with the Python code for processing the data and reproducing our results.

### Data availability

All datasets used in this study are publicly available. These include the PBMC 68k scRNA-seq dataset [48], the PBMC 10k scRNA-seq dataset [49], the human white adipose tissue data [30], the Xenium spatial transcriptomics dataset for human breast cancer [50], the bulk RNA-seq reference from Monaco et al. [51], and the FANTOM5 bulk RNA-seq reference atlas [52].

### Author contributions

XZ developed the method, conducted the analysis, wrote the manuscript. KALC and JM supervised the work, wrote and edited the manuscript.

## Acknowledgements

We would like to thank Dr Alexandr Garbali (University of Mebourne) for helpful discussions.

