## Supplemental material for "CellDiffusion: a generative model to annotate single-cell and spatial RNA-seq using bulk references"

### S1 CellDiffusion Model Implementation Details

#### S1.1 Theoretical Foundation of DDPMs

The DDPM framework involves two main processes: a fixed forward noising process and a learned reverse denoising process.

The forward process gradually introduces Gaussian noise to the initial data $x_0$ over a sequence of $T = 500$ discrete timesteps. This process is defined as a Markov chain that produces a series of progressively noisier samples $x_1, \ldots, x_T$, according to the conditional probability:

$$q(x_t|x_{t-1}) = \mathcal{N}(x_t; \sqrt{1-\beta_t}\, x_{t-1}, \beta_t I),$$

where $\mathcal{N}(x; \mu, \Sigma)$ denotes a Gaussian distribution with mean vector $\mu$ and covariance matrix $\Sigma$. The term $\{\beta_t\}_{t=1}^{T}$ is a predefined linear variance schedule, increasing from $\beta_1 = 0.0001$ to $\beta_T = 0.02$. $I$ is the identity matrix. This configuration ensures that after $T$ steps, the final sample $x_T$ converges to an isotropic Gaussian distribution $\mathcal{N}(\mathbf{0}, I)$.

The reverse process is a learned generative model designed to reverse the forward process. Starting from a sample of pure noise, $x_T \sim \mathcal{N}(\mathbf{0}, I)$, the model iteratively denoises the sample at each timestep to reconstruct a clean data point, $\hat{x}_0$.

#### S1.2 Model Architecture

The core of the CellDiffusion model is a U-Net, a convolutional neural network architecture renowned for its efficacy in image-to-image translation tasks [1]. Its symmetric encoder-decoder structure with skip connections is particularly well suited for reconstructing the detailed structure of the reshaped gene expression matrices.

The network architecture consists of a series of ResNet blocks, which utilise group normalisation (with 8 groups) and weight standardisation for stable training [2]. The encoder path progressively downsamples the input, with channel dimension multipliers of (1, 2, 4, 8) at successive resolutions. To capture contextual information effectively, we employ a dual attention system: a computationally efficient linear attention mechanism is used in the encoder and decoder blocks, while a full attention mechanism is applied at the bottleneck layer. The timestep $t$ is incorporated into the model via sinusoidal position embeddings, allowing the network to condition its output on the noise level.

#### S1.3 Model Training

The U-Net is trained to predict the noise component, $\varepsilon$, that was added to the input matrix $x_0$ at a given timestep $t$. To achieve this, the model is optimised by minimising the following objective function:

$$L_{\text{simple}} = \mathbb{E}_{t,x_0,\varepsilon}\left[\left\|\varepsilon - \varepsilon_\theta(x_t, t)\right\|^2\right],$$

where $\mathbb{E}$ denotes the expectation, $x_0$ is the original data matrix, $\varepsilon \sim \mathcal{N}(\mathbf{0}, I)$ is the sampled Gaussian noise, $x_t$ is the noisy sample at timestep $t$, and $\varepsilon_\theta$ represents the U-Net model parameterized by $\theta$. For increased robustness to outliers in the gene expression data, we implemented this objective using the Huber loss instead of the standard L2 norm.

Training was conducted for 300 epochs using the Adam optimiser [3] with a learning rate of $1 \times 10^{-3}$ and a batch size of 128. The model was implemented in Python (v3.8.12) using PyTorch (v1.11.0).

#### S1.4 Synthetic Virtual Cell Generation

Upon completion of training, the model generates synthetic gene expression profiles, termed "virtual cells", through its learned reverse denoising process. Generation begins by sampling an initial matrix from a standard Gaussian distribution ($x_T \sim \mathcal{N}(\mathbf{0}, I)$). This matrix is then iteratively denoised by the model over timesteps $t = T, T-1, \ldots, 1$ using the following update rule:

$$x_{t-1} = \frac{1}{\sqrt{\alpha_t}} \left( x_t - \frac{1-\alpha_t}{\sqrt{1-\bar{\alpha}_t}} \varepsilon_\theta(x_t, t) \right) + \sigma_t z,$$

where $\alpha_t = 1 - \beta_t$, $\bar{\alpha}_t = \prod_{i=1}^{t} \alpha_i$, $z \sim \mathcal{N}(\mathbf{0}, I)$ for $t > 1$ (with $z = 0$ for $t = 1$), and $\sigma_t$ is the noise variance.

The final output, $\hat{x}_0$, is a synthetic gene expression matrix representing a novel virtual cell. This procedure can be repeated to generate arbitrarily large cell populations for subsequent analyses. To ensure biological realism, a final post-processing step enforces nonnegativity by setting any negative expression values in $\hat{x}_0$ (usually very close to zero) to zero, which also enhances the natural sparsity of the data.

### S2 Details of datasets and data preprocessing

#### S2.1 Datasets

Our study utilised a combination of publicly available scRNA-seq, image-based spatial transcriptomics, and bulk RNA-seq datasets to evaluate model performance. The scRNA-seq and spatial transcriptomics datasets served as the query data, while the bulk RNA-seq datasets were used as the reference for training and annotation. A detailed summary of all datasets, their modality, and their specific roles in our benchmark and case study analyses is provided in Table S1.

**Table S1:** Overview of datasets used for model evaluation.

| Dataset Name | Modality | Role | Use Case | Ref. |
|---|---|---|---|---|
| PBMC 68k | scRNA-seq | Query | Benchmark | [4] |
| PBMC 10k | scRNA-seq | Query | Case Study | [5] |
| Adipose (Emont) | scRNA-seq | Query | Benchmark & Case Study | [6] |
| Xenium Breast CA | Spatial | Query | Benchmark & Case Study | [7] |
| Monaco | Bulk RNA-seq | Reference | Reference | [8] |
| FANTOM5 | Bulk RNA-seq | Reference | Reference | [9] |

#### S2.2 Single Cell RNA-seq Data Processing

For a given scRNA-seq dataset, the gene expression profile of each cell is first represented as a vector, $C = [g_1, g_2, \ldots, g_M]$, where $g_i$ is the expression count for the $i$-th gene and $M$ is the total number of genes.

We perform feature selection by identifying the top $N$ most highly variable genes expressed in at least three cells; by default, $N = 4,096$. Expression values for this gene subset are then normalised to a library size of $10^6$ (counts per million, CPM) and log-transformed via $\log_e(\text{CPM} + 1)$ to stabilise the variance.

To make the data compatible with the convolutional layers of the U-Net architecture, the $N$-dimensional gene vector is reshaped into a square matrix of size $n \times n$, where $n = \sqrt{N}$ (e.g., $64 \times 64$ for $N = 4,096$). If $N$ is not a perfect square, the vector is padded with zeros to complete the matrix dimensions. This final matrix is denoted as $x_0$ and serves as the input to the CellDiffusion model.

#### S2.3 Bulk RNA-seq Data Processing

Any bulk RNA-seq reference data used for downstream tasks undergoes an analogous preprocessing pipeline to ensure consistency. This includes selecting the set of $N$ genes that overlap with the processed scRNA-seq data, normalising to CPM, and applying a log transformation.

### S3 Details of Benchmark Study

To rigorously evaluate the performance of our method, we conducted a comprehensive benchmark against several state-of-the-art cell type annotation tools. The evaluation was performed on the query datasets described in Table S1.

#### S3.1 Methods compared

We compared our model against four established methods: SingleR, Seurat, CHETAH, and scVI/scANVI. Each tool was run using parameters recommended in its documentation.

- **SingleR:** We used the `SingleR` R package to perform reference-based annotation. Labels were assigned for each query cell by calling the main `SingleR()` function, using the query data as the `test` set and the bulk RNA-seq data as the `ref` set.
- **Seurat:** We employed the label transfer workflow from the `Seurat` R package. We first identified integration anchors between the reference and query datasets using the `FindTransferAnchors()` function, considering the top 30 principal components (`dims = 1:30`). Cell type labels were then transferred to the query data using the `TransferData()` function.
- **CHETAH:** We utilized the `CHETAH` R package for hierarchical classification. The `CHETAHclassifier()` function was used with the query single cell data as input and the bulk data as the reference. The confidence threshold was set to 0 (`thresh = 0`) to ensure all cells were assigned a label.
- **scVI/scANVI:** For this deep learning based approach, we first trained an scVI model on the integrated reference and query data. A scANVI model was then initialised from the trained scVI model using `SCANVI.from_scvi_model()`, designating the reference labels with the `labels_key` and query cells as an `unlabeled_category`. The scANVI model was subsequently trained for 100 epochs, and cell type predictions were generated using the `predict()` function.

#### S3.2 Performance Evaluation

The performance of each method was evaluated by comparing its predicted cell type labels against ground truth annotations. For each query dataset, these ground truth labels were derived from the expert manual annotations provided in the original publication, which served as the gold standard for our assessment.

To quantify performance, we calculated a suite of standard classification metrics using custom Python scripts leveraging the `scikit-learn` library. Recognising that single-cell transcriptomics datasets are often highly imbalanced—with some cell types being significantly rarer than others—we focused on metrics that are robust to such distributions. The metrics used for this benchmark included:

- **Accuracy:** The overall proportion of cells correctly annotated. While a useful general measure, it can be misleading in datasets with imbalanced cell type populations.
- **Macro-averaged Precision:** The unweighted mean of precision scores calculated for each cell type. Precision for a given cell type measures the proportion of true positives (correct annotation) among all instances predicted as that cell type ($TP/(TP+FP)$). This metric assesses the reliability of the annotations, indicating how likely a cell is to actually be of a certain type if it is predicted as such. Macro-averaging ensures that the performance on rare cell types contributes equally to the overall score as abundant types.
- **Macro-averaged Recall (Sensitivity):** The unweighted mean of recall scores for each cell type. Recall for a given class measures the proportion of all actual cells of that type that were correctly identified ($TP/(TP+FN)$). This metric evaluates the model's ability to find all instances of a particular cell type and is particularly important for assessing the detection of rare cell populations.
- **Macro F1-Score:** The unweighted average of the F1-scores for each cell type. The F1-score is the harmonic mean of precision and recall, providing a single metric that balances both concerns. It is especially suitable for imbalanced datasets as it treats all classes equally, regardless of their size.
- **Cohen's Kappa:** A statistic that measures the agreement between predicted and true labels while correcting for the probability of agreement occurring by chance. It provides

a more robust measure than simple accuracy, especially when class distributions are imbalanced.

### S4 Details of Downstream Analyses for Case Studies

#### S4.1 Downstream Analyses for the Monocyte Subtypes Case Study

All downstream analyses for the monocyte case study were performed in Python using the `scanpy` package [10].

**Data Subsetting and Preprocessing** First, cells annotated as Monocyte by CellDiffusion were computationally isolated from the complete Peripheral Blood Mononuclear Cell (PBMC) dataset. This monocyte-specific subset was then independently preprocessed. Gene counts were normalised to a library size of 10,000 per cell (`scanpy.pp.normalize_total`), and highly variable genes (HVGs) were identified (`scanpy.pp.highly_variable_genes`). The expression data for these HVGs was then scaled to have zero mean and unit variance, with values clipped at a maximum of 10 (`scanpy.pp.scale`).

**Dimensionality Reduction and Visualisation** Principal Component Analysis (PCA) was performed on the HVGs. The top 40 principal components were used to construct a k-nearest neighbour graph (k=10; `scanpy.pp.neighbors`). This graph was then used to compute the Uniform Manifold Approximation and Projection (UMAP) embedding for visualisation of the cellular landscape.

**Subtype Quantification and Marker Gene Validation** The relative proportions of monocyte subtypes (classical, intermediate, and nonclassical), as annotated by CellDiffusion, were quantified. To validate these assigned identities, we visualised the expression of canonical marker genes `CD14` and `FCGR3A` on the UMAP embedding and using violin plots.

**Differential Gene Expression Analysis** To uncover the transcriptional signatures defining each monocyte subtype, differential gene expression (DGE) analysis was performed using the Wilcoxon rank sum test (`scanpy.tl.rank_genes_groups`). This procedure identified genes significantly upregulated in each subtype relative to all other subtypes.

**Functional Enrichment Analysis** To interpret the biological functions associated with each monocyte subtype, we performed functional enrichment analysis on the DGE results. For each subtype, genes with $log_2FC > 1.0$ and adjusted $p - value < 0.05$ were selected. This gene set was analysed using the `gprofiler` Python package (v1.0.0) to query the Gene Ontology, KEGG, and Reactome pathway databases. The analysis was set to *Homo sapiens*, with a significance threshold of $FDR < 0.05$ [11]. For each subtype with at least five significant genes, the top 10 most enriched terms were retained for interpretation.

**Pseudotime and Trajectory Inference** To investigate the differentiation trajectories of monocyte subtypes, we performed pseudotime analysis using diffusion pseudotime (DPT). A neighbourhood graph was constructed on the 40-dimensional PCA space (k=30 neighbours), and cellular clusters were identified using the Leiden algorithm (resolution=3.0). Outlier clusters containing fewer than 10 cells were removed.

The root cell for DPT calculation was selected from Leiden cluster 17, which was identified based on high expression of early monocyte markers (`LYZ`, `S100A9`). Specifically, the cell closest to the cluster centroid in UMAP space was chosen as the root to ensure a representative starting point for the trajectory. The diffusion map was computed using the first 10 diffusion components. This approach ordered cells along a continuous trajectory, modelling the putative progression from classical monocytes through intermediate states to nonclassical monocytes. To validate the trajectory, the expression of canonical markers (`CD14` and `FCGR3A`) was examined along the pseudotime axis. Gene expression trends were visualised by plotting smoothed expression values (locally weighted scatterplot smoothing; LOWESS, factor=0.3) against pseudotime.

#### S4.2 Downstream Analyses for the Adipose Tissue Case Study

This case study focused on the myeloid cell populations from the single-cell transcriptome data of Emont et al. [6].

**Data Preprocessing and Dimensionality Reduction** The raw data were processed using a pipeline similar to the monocyte analysis, including library size normalisation, log transformation, and HVGs identification. PCA was performed on the HVG matrix, and the top 30 principal components were used to construct a k-NN graph, which served as the basis for UMAP embedding.

**Cell Type Annotation** For comparison, we employed two strategies for cell type annotation. First, the auto machine learning classifier utilised a multilayer perceptron (MLP) model trained on the FANTOM5 bulk RNA-seq dataset. This model was applied to pseudobulk profiles generated by the CellDiffusion framework to classify cells into major myeloid lineages. Second, as an independent approach, the query dataset was mapped to the Azimuth 'Human Bone Marrow' reference map to obtain a separate set of cell labels [12–14].

**Marker Gene Identification and Functional Analysis** Marker genes for each annotated cell type were identified via differential gene expression (DGE) analysis using the Wilcoxon rank sum test (`scanpy.tl.rank_genes_groups`). The resulting lists of significantly upregulated genes were then subjected to functional enrichment analysis using the `gseapy` Python package against the Gene Ontology (GO) Biological Process 2021 database to infer the biological functions of each cell type [15].

**Trajectory and Pseudotime Analysis** To investigate differentiation dynamics within the myeloid compartment, we performed trajectory inference using Partition-based graph abstraction (PAGA) in conjunction with Diffusion Pseudotime (DPT). The PAGA graph was first constructed to map the connectivity between cell clusters. A root for the trajectory was defined within the monocyte population based on biological priors, from which pseudotime values were computed. This analysis allowed for the ordering of cells along a continuous monocyte to macrophage differentiation path and the visualisation of gene expression changes, such as for key markers `FABP4` and `NAMPT`, along this trajectory.

### S4.3 Downstream Analyses for the Image-based Spatial Transcriptomics Case Study

To investigate the spatial organisation of the tumour microenvironment (TME), we conducted downstream analyses on the annotated Xenium spatial transcriptomics data from breast cancer [7]. All analyses utilized custom Python scripts leveraging the `scanpy` (v1.9.1) and `squidpy` (v1.2.3) packages [10, 16].

**Cell Type Annotation and Data Subsetting** The Xenium breast cancer dataset was first annotated using the CellDiffusion framework with the Monaco bulk RNA-seq atlas as a reference. From the annotated data, we computationally isolated the tumour region based on annotations provided in the metadata of the original publication. For proximity analyses, a subset of immune cells comprising $CD4^+$ T cells, $CD8^+$ T cells, and B cells was selected.

**Statistical Analysis of Immune Tumour Proximity** To quantify the spatial relationship between immune and tumour cells, we performed a proximity analysis. For each immune cell of interest, we calculated the Euclidean distance to the nearest tumour cell based on their spatial coordinates. An immune cell was defined as being in proximity to a tumour if this distance was less than or equal to 200µm.

The statistical significance of this colocalization was assessed with a permutation test. To generate a null distribution for each immune subtype, we performed 1,000 permutations where the observed number of immune cells was randomly repositioned within the tissue boundaries. For each permutation, we calculated the proportion of randomly placed cells falling within the 200 µm proximity threshold. The p-value was determined as the fraction of permutations where the resulting proximity proportion was equal to or greater than the observed proportion. A $p - value < 0.05$ was considered statistically significant, indicating a nonrandom spatial association.

**Immune Cell Enrichment and Density Analysis** We performed a spatial enrichment analysis to determine if specific immune populations were enriched within the annotated tumour region. This analysis compared the observed count of each immune subtype within the tumour boundary to the count expected under a null hypothesis of random spatial distribution. Additionally, local cell densities were calculated to characterise immune ”hot” and ”cold” spots across the tissue.

**T cell Subtype Analysis** To resolve T cell heterogeneity, we performed a subclustering analysis on all cells annotated as T cells. Subtype identities were assigned by referencing the Monaco dataset. The spatial coordinates of the resulting T cell subtypes were then visualised to map their distribution and localisation within the TME.

[11] Uku Raudvere, Liis Kolberg, Ivan Kuzmin, Tambet Arak, Priit Adler, Hedi Peterson, and Jaak Vilo. g: Profiler: a web server for functional enrichment analysis and conversions of gene lists (2019 update). *Nucleic acids research*, 47(W1):W191–W198, 2019.

[12] Yuhan Hao, Stephanie Hao, Erica Andersen-Nissen, William M Mauck, Shiwei Zheng, Andrew Butler, Maddie J Lee, Aaron J Wilk, Charlotte Darby, Michael Zager, et al. Integrated analysis of multimodal single-cell data. *Cell*, 184(13):3573–3587, 2021.

[13] Karolyn A Oetjen, Katherine E Lindblad, Meghali Goswami, Gege Gui, Pradeep K Dagur,

Catherine Lai, Laura W Dillon, J Philip McCoy, and Christopher S Hourigan. Human bone marrow assessment by single-cell rna sequencing, mass cytometry, and flow cytometry. *JCI insight*, 3(23):e124928, 2018.

[14] Jeffrey M Granja, Sandy Klemm, Lisa M McGinnis, Arwa S Kathiria, Anja Mezger, M Ryan Corces, Benjamin Parks, Eric Gars, Michaela Liedtke, Grace XY Zheng, et al. Single-cell multiomic analysis identifies regulatory programs in mixed-phenotype acute leukemia. *Nature biotechnology*, 37(12):1458–1465, 2019.

[15] Zhuoqing Fang, Xinyuan Liu, and Gary Peltz. Gseapy: a comprehensive package for performing gene set enrichment analysis in python. *Bioinformatics*, 39(1):btac757, 2023.

[16] Giovanni Palla, Hannah Spitzer, Michal Klein, David Fischer, Anna Christina Schaar, Louis Benedikt Kuemmerle, Sergei Rybakov, Ignacio L Ibarra, Olle Holmberg, Isaac Virshup, et al. Squidpy: a scalable framework for spatial omics analysis. *Nature methods*, 19(2): 171–178, 2022.
